# NrnA is a linear dinucleotide phosphodiesterase with limited function in cyclic dinucleotide metabolism in *Listeria monocytogenes*

**DOI:** 10.1101/2021.04.21.440870

**Authors:** Aaron R. Gall, Cheta Siletti, TuAnh N. Huynh

## Abstract

*Listeria monocytogenes* produces both c-di-AMP and c-di-GMP to mediate many important cellular processes, but the levels of both nucleotides must be regulated. C-di-AMP accumulation attenuates virulence and diminishes stress response, and c-di-GMP accumulation impairs bacterial motility. An important regulatory mechanism to maintain c-di-AMP and c-di-GMP homeostasis is to hydrolyze them to the linear dinucleotides pApA and pGpG, respectively, but the fates of these hydrolytic products have not been examined in *L. monocytogenes*. We found that NrnA, a stand-alone DHH-DHHA1 phosphodiesterase, has a broad substrate range, but with a strong preference for linear dinucleotides over cyclic dinucleotides. Although NrnA exhibited detectable cyclic dinucleotide hydrolytic activities in vitro, NrnA had negligible effects on their levels in the bacterial cell, even in the absence of the c-di-AMP phosphodiesterases PdeA and PgpH. The Δ*nrnA* mutant had a mammalian cell infection defect that was fully restored by *E. coli* Orn. Together, our data indicate that *L. monocytogenes* NrnA is functionally orthologous to Orn, and its preferred physiological substrates are most likely linear dinucleotides. Furthermore, our findings revealed that, unlike some other c-di-AMP and c-di-GMP-producing bacteria, *L. monocytogenes* does not employ their hydrolytic products to regulate their phosphodiesterases, at least at the pApA and pGpG levels in the Δ*nrnA* mutant. Finally, the Δ*nrnA* infection defect was overcome by constitutive activation of PrfA, the master virulence regulator, suggesting that accumulated linear dinucleotides might inhibit the expression, stability, or function of PrfA-regulated virulence factors.

**IMPORTANCE:** *Listeria monocytogenes* produces both c-di-AMP and c-di-GMP, and encodes specific phosphodiesterases that degrade them into pApA and pGpG, respectively, but the metabolism of these products has not been characterized in this bacterium. We found that *L. monocytogenes* harbors an NrnA homolog that degrades a broad range of nucleotides, but exhibits a strong biochemical and physiological preference for linear dinucleotides, including pApA and pGpG. Unlike in some other bacteria, these oligoribonucleotides do not appear to interfere with cyclic dinucleotide hydrolysis. The absence of NrnA is well tolerated by *L. monocytogenes* in broth cultures but impairs its ability to infect mammalian cells. These findings indicate a separation of cyclic dinucleotide signaling and oligoribonucleotide metabolism in *L. monocytogenes*.

## INTRODUCTION

C-di-AMP and c-di-GMP are among the most widespread cyclic dinucleotide second messengers in bacteria. Together, they are produced by all bacterial phyla, but their phylogenetic distributions do not entirely overlap (1). These nucleotides mediate many important aspects of bacterial physiology and pathogenesis, and c-di-AMP is also essential for the growth of many Firmicute species in rich laboratory media and in the infected hosts (2, 3). However, cyclic dinucleotide levels must be regulated to avoid toxic accumulation, in part through hydrolysis. Depending on the phosphodiesterases, c-di-AMP and c-di-GMP may be degraded into pApA and pGpG, respectively, or into AMP and GMP (4, 5).

In addition to cyclic dinucleotide hydrolysis, pApA and pGpG are also products of RNA metabolism. During bacterial growth, mRNA decay, abortive transcription initiation, and transcription elongation all generate RNA fragments that are cleaved by various RNases into oligoribonucleotides of 2-5 nucleotides, including pApA and pGpG (6, 7). These oligoribonucleotides, also called nanoRNAs, must be further degraded by nanoRNases to avoid detrimental consequences on bacterial growth and physiology. Indeed, the accumulation of oligoribonucleotides is lethal in *Escherichia coli* (8). Although the mechanisms for their toxic effects are not fully understood, nanoRNAs of 2-4 nucleotides can prime and shift transcription start sites at a large number of promoters in *Pseudomonas aeruginosa* (9). These oligoribonucleotides likely also regulate CRISPR-associated gene expression in *Mycobacterium* sp., but do not globally alter the transcriptional profile (10).

For some c-di-AMP and c-di-GMP-producing bacteria, there appears to be crosstalk between cyclic dinucleotide signaling and oligoribonucleotide metabolism. In *Staphylococcus aureus*, pApA inhibits the c-di-AMP phosphodiesterase activity of GdpP (also called PdeA); and in *Pseudomonas aeruginosa*, pGpG inhibits the c-di-GMP phosphodiesterase activity of EAL-domain protein RocR (11–13). A subset of nanoRNases, including Orn, NrnA, NrnB, and NrnC can readily degrade pGpG (14). Indeed, Orn has a strong preference for linear dinucleotides in bacterial cells over longer substrates (15). Accordingly, a *P. aeruginosa* Δ*orn* mutant, which is severely diminished for pGpG degradation, also accumulates c-di-GMP (11, 12). Furthermore, several NrnA homologs, also called DhhP, can hydrolyze c-di-AMP and c-di-GMP (5). NrnA is the only identified hydrolase for both c-di-AMP and oligoribonucleotides in *Mycobacterium* sp. (16, 17).

*Listeria monocytogenes* is a Gram-positive Firmicute bacterium that produces both c-di-AMP and c-di-GMP. C-di-AMP directly binds several protein targets and mediates many important cellular processes, such as central metabolism, osmolyte transport, and cell wall integrity (18– 20). C-di-AMP homeostasis is critical to *L. monocytogenes* growth, stress response, and pathogenesis. Whereas c-di-AMP is essential for growth in standard laboratory media and during infection, its accumulation also greatly attenuates virulence (21, 22). Similarly, c-di-GMP levels must be regulated, since its build-up promotes *L. monocytogenes* cell aggregation and impairs mammalian cell invasion (23, 24). While substantial progress has been made on the molecular targets and regulatory mechanisms of cyclic dinucleotide signaling in *L. monocytogenes*, much less is known about the fate of their hydrolytic products, pApA and pGpG.

Among the nanoRNases that have been identified in other bacteria, *L. monocytogenes* encodes only NrnA, which is a stand-alone DHH-DHHA1 phosphodiesterase. Here, we found that *L. monocytogenes* NrnA is a promiscuous enzyme in vitro, but has a strong preference for linear dinucleotides over cyclic dinucleotides and pAp. Consistent with biochemical data, NrnA appears to have minimal hydrolytic activities towards c-di-AMP and c-di-GMP in the bacterial cell, both in broth cultures and infected mammalian cells. The Δ*nrnA* mutant has a biofilm formation and infection defect, most likely due to the accumulation of linear dinucleotides.

## RESULTS

### *Lm* NrnA is a promiscuous enzyme in vitro, with activities towards pAp, cyclic- and linear dinucleotides

Like other bacteria in the Firmicutes phylum, *L. monocytogenes* does not encode an Orn ortholog. Instead, it harbors a single NrnA homolog, encoded by *lmo1575*, with a stand-alone DHH-DHHA1 catalytic domain that belongs to the DHH phosphodiesterase superfamily. *Lm* NrnA is highly similar in amino acid sequence to *B. subtilis* NrnA, with 54% sequence identity and 63% sequence similarity, and the two sequences are co-linear (**Fig. S1**). The recombinant *Lm* NrnA-6xHis protein exhibited a robust phosphodiesterase activity towards the model substrate bis-*p*-nitrophenylphosphate (bis-*p*NPP). The enzymatic activity was optimal at pH 8.0 and dependent on Mn^2+^, consistent with several DHH-DHHA1 phosphodiesterases (25–28) (**Fig. S2**).

*L. monocytogenes* produces both c-di-AMP and c-di-GMP, which are hydrolyzed into pApA and pGpG, respectively. To understand *Lm* NrnA functions in cyclic dinucleotide metabolism, we first examined its enzymatic activities towards these nucleotide substrates. Under our assay conditions, *Lm* NrnA was active against all these cyclic and linear dinucleotides (**Fig.1** and **Fig. S3**). Although the affinities for all compounds were in the same order of magnitude, catalytic efficiencies (*k*_cat_/K_m_) for pApA and pGpG were 200 – 300 fold higher than those for c-di-AMP and c-di-GMP (**Table 1**). Using pApG as a proxy for other oligoribonucleotides unrelated to cyclic dinucleotide signaling in *L. monocytogenes*, we found that NrnA also robustly hydrolyzed this substrate, at similar affinity and efficiency to pApA and pGpG (**Fig.1** and **Table 1**). Finally, because the *B. subtilis* NrnA homolog also degrades 3’-phosphoadenosine-5’-phosphate (pAp) (29), we tested *Lm* NrnA towards this substrate, and observed a readily detectable activity, albeit at a much lower catalytic efficiency than that for linear dinucleotides (**Table 1**).

**Figure 1:**
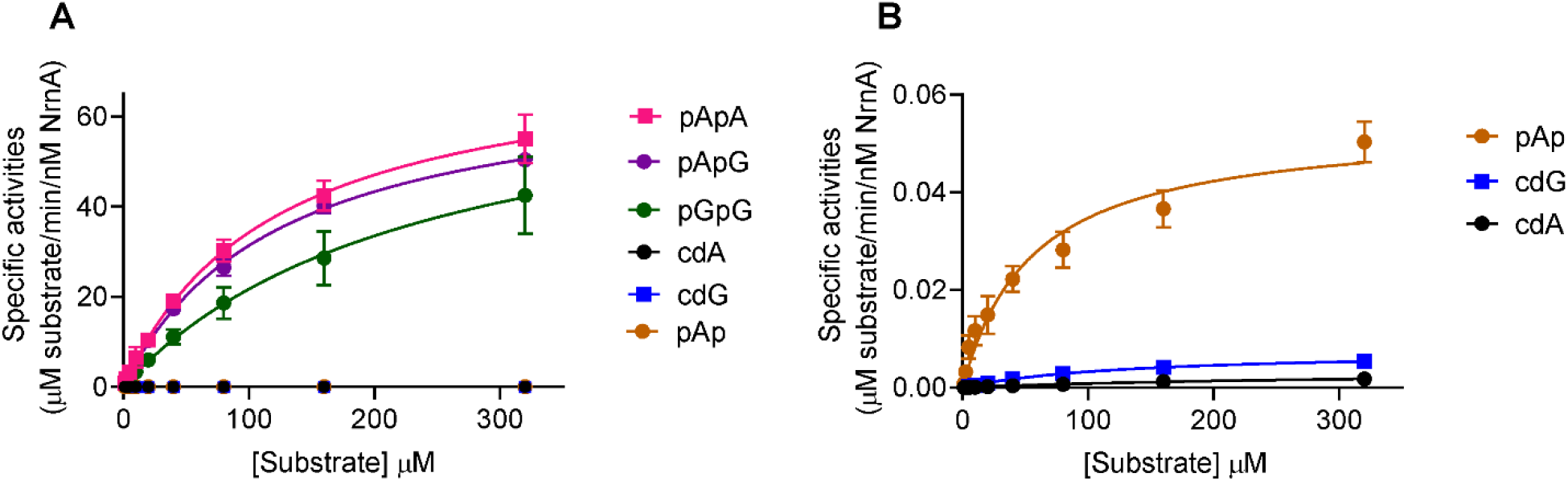
*Lm* NrnA is a promiscuous phosphodiesterase/phosphatase with a strong preference for linear dinucleotides. **A** – Enzymatic activities towards all tested substrates. Abbreviations: cdA = c-di-AMP, cdG = c-di-GMP. Reactions were stopped and quantified for substrates and products by HPLC, using a standard curve for each compound. Specific activities are average of at least three independent experiments. **B** – NrnA activities towards pAp, c-di-AMP, and c-di-GMP, replotted from (A) on a different y-axis scale.

**Table 1:** *Lm* NrnA kinetic constants, generated by fitting data from Fig. 1 to the Michaelis-Menten equation in GraphPad Prism version 9.0.1

| Compound | $k_{\text{cat}}$ ($\text{min}^{-1}$) | $K_{\text{m}}$ ($\mu\text{M}$) | $k_{\text{cat}}/K_{\text{m}}$ ($\mu\text{M}^{-1}\text{min}^{-1}$) |
| --- | --- | --- | --- |
| pApA | $75 \pm 3$ | $119 \pm 11$ | 0.639 |
| pGpG | $74 \pm 12$ | $239 \pm 74$ | 0.307 |
| pApG | $70 \pm 2$ | $124 \pm 7$ | 0.566 |
| pAp | $0.054 \pm 0.003$ | $55.4 \pm 9.7$ | $97 \times 10^{-5}$ |
| c-di-AMP | $0.004 \pm 0.0004$ | $338 \pm 62$ | $1 \times 10^{-5}$ |
| c-di-GMP | $0.007 \pm 0.0004$ | $133 \pm 16$ | $5 \times 10^{-5}$ |

The DHH motif is invariant among DHH superfamily proteins, and is indispensable for NrnA catalytic activity, since the middle His residue of this motif coordinates divalent metals for hydrolysis (25, 27). Consistent with this requirement, *Lm* NrnA activity was completely abolished when the DHH motif was substituted with AAA residues (**Fig. S2A**).

In addition to NrnA, *B. subtilis* harbors NrnB and YhaM, both of which have been tested for hydrolytic activities towards oligoribonucleotides (30). *L. monocytogenes* does not harbor NrnB, but has a YhaM homolog, encoded by *lmo2220. Lm* YhaM, purified as a maltose-binding protein fusion, exhibited phosphodiesterase activity towards bis-*p*NPP (**Fig. S4**), but not any of the nucleotide substrates even at high enzyme concentrations and prolonged incubation (data not shown).

### *Lm* NrnA does not hydrolyze c-di-AMP and c-di-GMP in *L. monocytogenes* broth cultures or during infection

In some c-di-AMP and c-di-GMP producing bacteria, a feedback regulatory loop has been reported between cyclic dinucleotide levels and their hydrolytic products (11–13). Thus, we sought to determine whether *Lm* NrnA was involved in the regulation of cyclic dinucleotide homeostasis. *L. monocytogenes* synthesizes c-di-AMP by the diadenylate cyclase DacA, and degrades it by at least two phosphodiesterases, PdeA and PgpH (21, 22) (**Fig. 2A**). In the WT, *dacA-*depleted, and *pdeA pgpH* mutant backgrounds, *nrnA* deletion had no effects on c-di-AMP levels (**Fig. 2B**). Additionally, in the *pdeA pgpH* mutant, which accumulates c-di-AMP four-fold compared to WT, *nrnA* over-expression did not reduce c-di-AMP levels. Altogether, these data indicate that NrnA has a minor or negligible role in c-di-AMP hydrolysis in *L. monocytogenes*. Our data also suggest that, in the absence of NrnA, the accumulated pApA did not significantly inhibit c-di-AMP hydrolysis.

**Figure 2:**
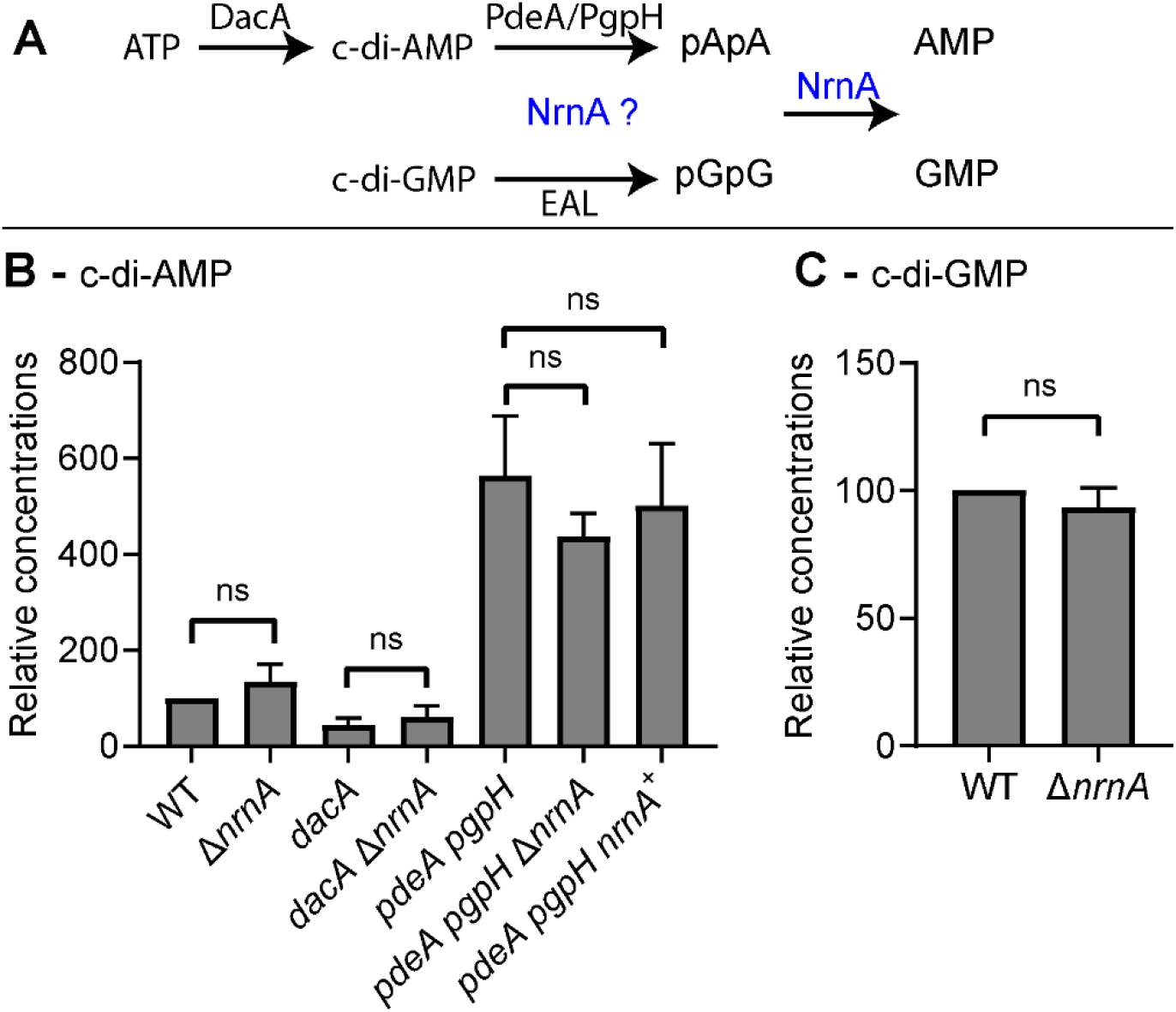
*Lm* NrnA has minimal c-di-AMP and c-di-GMP phosphodiesterase activity in *L. monocytogenes*. **A** – Schematic diagram of c-di-AMP and c-di-GMP metabolism in *L. monocytogenes*. **B** – Mid-exponential phase cultures grown in BHI broth were quantified for c-di-AMP levels by LC-MS/MS, with C^13^N^15^-c-di-AMP as an internal standard. In each experiment, c-di-AMP level of each strain was normalized to the WT level. Cytoplasmic c-di-AMP concentration for the WT strain was 7.6 ± 0.7 μM (standard deviation). **C** – C-di-GMP was quantified by LC-MS/MS similar to B. Cytoplasmic c-di-GMP concentrations for the WT strain was 12 ± 7 μM (standard deviation). The *pdeA pgpH* null allele mutants were either Δ*pdeA* Δ*pgpH* or *pdeA*::Tn917 Δ*pgpH* (Table S1). Strain *dacA* is a genetic depletion of the *dacA* gene. Data in B and C are average of at least three independent experiments. Error bars represent standard deviations. Statistical analyses were performed by paired t-tests comparing strains from each experiment, ns: non-significant.

*L. monocytogenes* is an intracellular pathogen that can invade many mammalian cell types and robustly replicate in the infected cell cytosol (31). C-di-AMP secreted by *L. monocytogenes* in the infected cell cytosol robustly activates type I interferon response (type I IFN) (32). Thus, we quantified type I IFN to assess c-di-AMP synthesis and secretion in infected cells (**Fig. 3**). As expected, the Δ*hly* mutant, which is trapped in the phagosome, did not induce type I IFN; whereas the *tetR*::Tn917 mutant, which hyper-secretes c-di-AMP, induced a robust response (32). The WT and Δ*nrnA* mutant elicited similar type I IFN responses, suggesting that these two strains had similar c-di-AMP levels. As also previously shown, the *pdeA pgpH* mutant hyper-induced type I IFN, and we found this response to be unaffected by *nrnA* deletion or over-expression (**Fig. 3**). Thus, these assays indicate that NrnA did not significantly degrade c-di-AMP in cytosolic *L. monocytogenes*.

**Figure 3:**
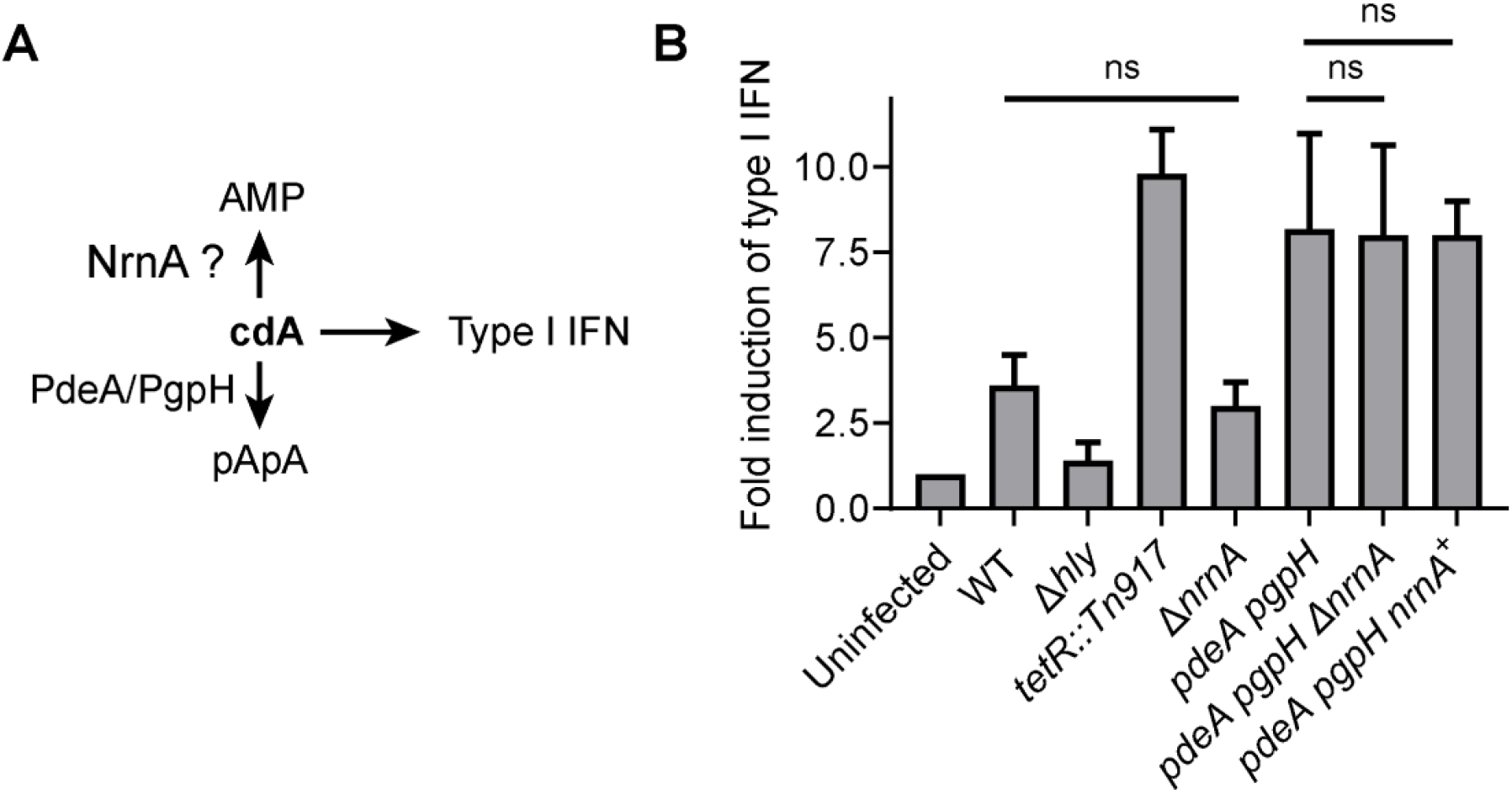
*Lm* NrnA has insignificant c-di-AMP phosphodiesterase activity during infection. **A** – *L. monocytogenes* induces type I interferon response (Type I IFN) by c-di-AMP. **B** – Type I IFN was quantified by ISRE assays. Bone marrow-derived macrophages were infected with *L. monocytogenes*, and cell supernatants were incubated with ISRE-L929 cells, and type I IFN was quantified based on bioluminescence, corrected for background level induced by reagents and media. Type I IFN responses were normalized to the level detected in uninfected cells in each experiment. Strains Δ*hly* and *tetR*::Tn917 were examined as negative and positive controls, respectively. The *pdeA pgpH* null allele mutants were either Δ*pdeA* Δ*pgpH* or *pdeA*::Tn917 Δ*pgpH* (Table S1). Data are average of three independent experiments. Error bars represent standard deviations. Statistical analyses were performed by paired t-test for responses in each experiment.

In addition to c-di-AMP, *L. monocytogenes* produces c-di-GMP, which is degraded into pGpG by three EAL-domain phosphodiesterases (23) (**Fig. 2A**). We found cytoplasmic c-di-GMP and c-di-AMP concentrations to be comparable in BHI-grown *L. monocytogenes*, at approximately ∼ 7 μM. The Δ*nrnA* mutant had a similar c-di-GMP level to WT, again indicating that NrnA had a minor role in cyclic dinucleotide hydrolysis in *L. monocytogenes* (**Fig. 2C**).

### The *L. monocytogenes* Δ*nrnA* mutant is diminished for biofilm formation

The *L. monocytogenes* Δ*nrnA* mutant had no growth defect in BHI broth (a rich medium) or *Listeria* Synthetic Medium (LSM, a chemically defined medium) (19). We also examined this mutant for growth under NaCl stress (varying from 0 – 6%) and antibiotics targeting the bacterial cell wall (β-lactams), transcription (rifampicin), and translation (chloramphenicol). The Δ*nrnA* mutant was indistinguishable from WT under these culture conditions (data not shown).

The *P. aeruginosa* Δ*orn* mutant is increased for biofilm formation due to an accumulation of c-di-GMP (11, 12, 14). Because the *L. monocytogenes* Δ*nrnA* mutant had a similar c-di-GMP level to WT, we did not anticipate any change in biofilm formation. Interestingly, we found the Δ*nrnA* strain to be diminished for biofilm formation in modified LSM, even though its total cell growth had the same density as WT (**Fig. 4A**).

**Figure 4:**
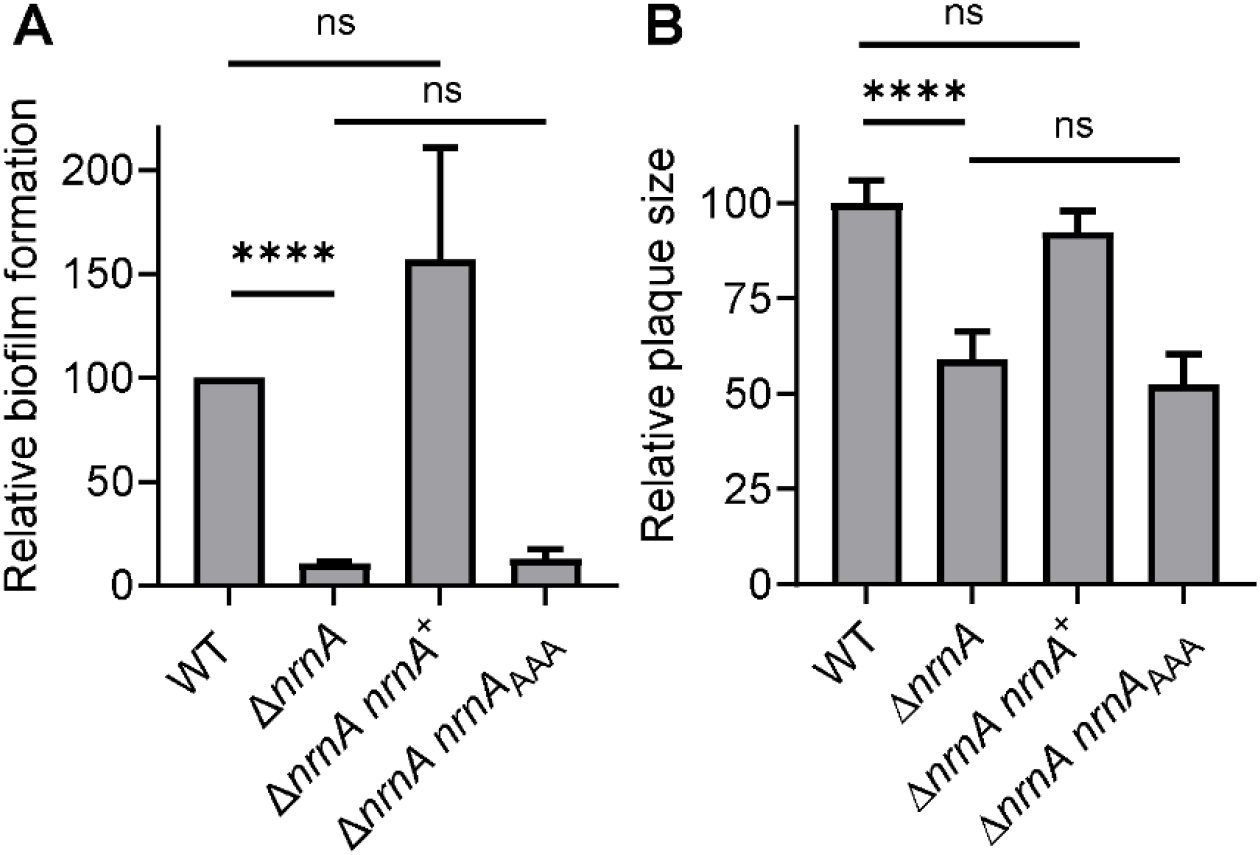
The *L. monocytogenes* Δ*nrnA* mutant is defective for biofilm formation and mammalian cell infection, due to the loss of NrnA enzymatic activity. **A** – Biofilm formation assays were performed for bacterial cultures grown in modified *Listeria* Synthetic Medium for 40 hours under static incubation at 30°C. Biofilm was quantified by staining with 0.3% crystal violet solution and read for absorbance at 595nm, and was normalized to the total culture density (OD 600nm) after thorough mixing of biofilm and planktonic growth biomass. **B** – Plaque formation at 4-6 days post infection of L2 fibroblasts. Plaque sizes were quantified by ImageJ. In each experiment, the average plaque size by each strain was normalized to the average WT plaque size. For both A and B, data are average of at least three independent experiments. Error bars represent standard deviations. Statistical analyses were performed by paired t-tests for strains from each experiment, ns, non-significant; ****, P < 0.0001.

### The *L. monocytogenes ΔnrnA* mutant has an infection defect, likely due to impaired PrfA-regulated virulence factors

As an intracellular pathogen, *L. monocytogenes* replicates in the infected cell cytosol and spreads to neighboring cells to avoid innate immune killing (31). Following host cell entry, the bacterium is enclosed in a phagosomal vacuole, and must escape into the cell cytosol through the activities of listeriolysin O (LLO, encoded by the *hly* gene). Cytosolic bacteria can rapidly replicate and spread to neighboring cells, mediated by ActA, which recruits and polymerizes host cell actin (31). Together with several virulence factors, LLO and ActA are transcriptionally regulated by the master virulence regulator PrfA, which is transcriptionally and allosterically activated during infection (33).

The *L. monocytogenes* intracellular lifecycle can be assessed by plaque formation upon infection of murine fibroblasts (L2 cells). As also previously reported, the Δ*nrnA* mutant exhibited a significant plaque formation defect, with an average plaque size at ∼ 60% of that formed by WT (**Fig. 4B**) (34). Consistent with this plaque defect, an *nrnA*::Himar1 mutant has also been shown to be diminished for virulence in the mouse infection model (34).

The Δ*nrnA* plaque formation defect might be due to diminished intracellular growth or cell-to-cell spread. To assess intracellular growth, we monitored the Δ*nrnA* mutant for replication in bone marrow-derived macrophages, and found that it was indistinguishable from the WT strain (**Fig. 5A**). We next evaluated cell-to-cell spread by examining *actA* expression. Because PrfA regulon expression is very weak in BHI broth, we exposed *L. monocytogenes* cultures to LB broth supplemented with glucose-1-phosphate and charcoal, a condition that robustly activates PrfA through unknown mechanisms (35, 36). Both the WT and Δ*nrnA* strains robustly upregulated *actA* expression under this condition (**Fig. 5B**). Of note, *hly* gene expression was also similar for both strains (**Fig. 5B**), despite the observation that the Δ*nrnA* mutant is diminished for β-hemolysis (data not shown and (34)).

**Figure 5:**
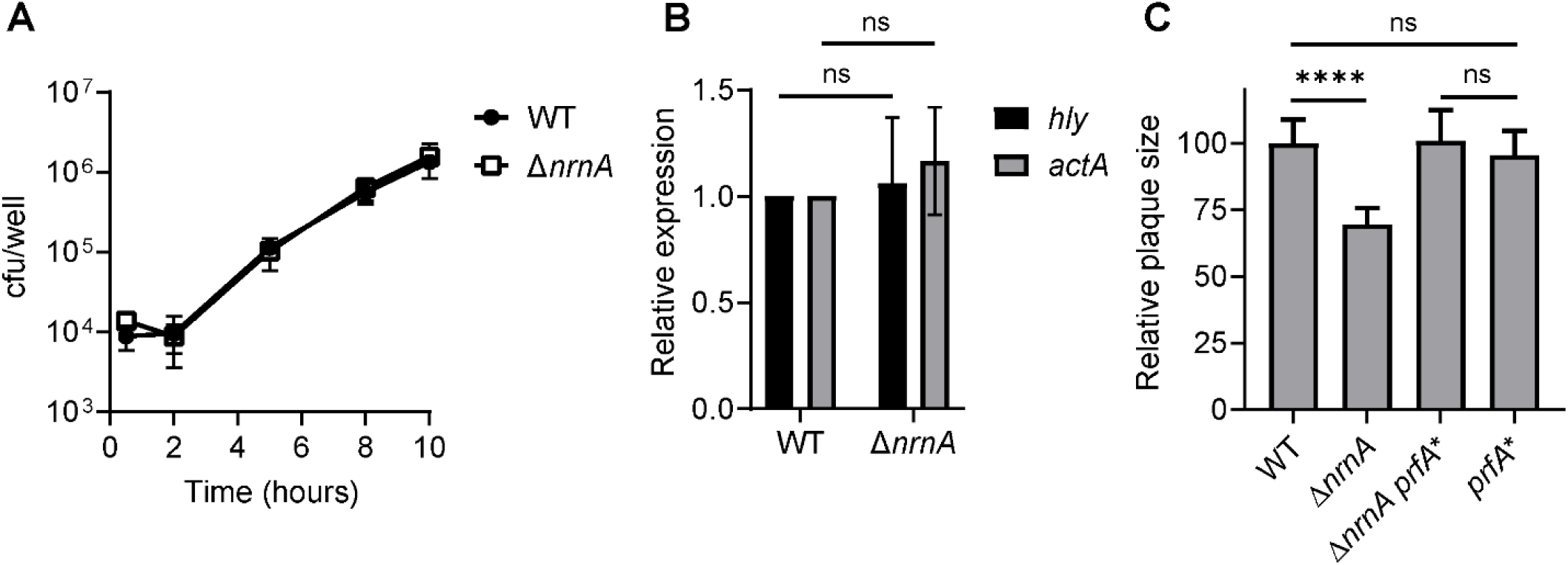
The *L. monocytogenes* Δ*nrnA* mutant has no defect for intracellular growth in infected macrophages, or expression of *actA*, which mediates cell-to-cell spread. **A** – Intracellular growth curves during infection of bone marrow-derived macrophages. **B** – Expression of *actA* and *hly* genes under PrfA-activating condition (exposure to glucose-1-phosphate and activated charcoal). Gene expression was normalized to that of the housekeeping gene *rplD*, and represented as relative to WT level. **C** – Plaque formation assays, performed as described in Fig. 4. PrfA* denotes PrfA G145S mutant, which is constitutively active. Data in A and B are average of at least two independent experiments. Data in C are average of at least three experiments. Error bars represent standard deviations. Statistical analyses were performed by paired t-tests for strains in each experiment: ns, non-significant; ****, P < 0.0001.

In addition to *hly* and *actA*, PrfA also upregulates several other virulence genes essential for the intracellular lifecycle (31). Although PrfA expression and activity are controlled by multiple mechanisms, there are several PrfA variants that are constitutively active, independent of these regulations (37, 38). The Δ*nrnA* plaque defect was completely rescued upon allelic replacement of wild-type PrfA with PrfA G145S (denoted as PrfA*), which is constitutively active (**Fig. 5C**). This indicates that the Δ*nrnA* strain was diminished for the expression, stability, or activity of virulence factors within the PrfA regulon.

### The Δ*nrnA* mutant phenotypes are likely due to oligoribonucleotide accumulation

For both biofilm formation in modified LSM and plaque formation during L2 cell infection, the Δ*nrnA* phenotypes were restored upon complementation with the WT *nrnA*^+^ allele, but not the *nrnA*_*AAA*_ (DHH → AAA) mutant allele, indicating that the biological function of NrnA is associated with its enzymatic activity (**Fig. 4**). Among different substrates tested in vitro, our analyses revealed that NrnA likely does not degrade c-di-AMP and c-di-GMP in *L. monocytogenes*. Thus, we sought to examine the physiological relevance of oligoribonucleotides and pAp. In *E. coli, M. tuberculosis*, and *B. subtilis*, pAp is a by-product of sulfate assimilation, encoded by *cysH, cysDNC* or analogous genes and operons (39). *L. monocytogenes* does not harbor any of these genes, but we could not exclude the possibility that it might still produce pAp from other unidentified pathways. Several bacteria encode CysQ that specifically degrades pAp, whereas Orn is specific for oligoribonucleotides (15, 39, 40). To distinguish these substrates in the bacterial cells, we complemented the Δ*nrnA* mutant with the *orn* or *cysQ* gene from *E. coli*. In both biofilm and plaque assays, Orn completely restored Δ*nrnA* defects, whereas CysQ had no detectable effect (**Fig. 6**). These data suggest that *Lm* NrnA is functionally analogous to Orn. Thus, the Δ*nrnA* phenotypes were caused by an accumulation of oligoribonucleotides, most likely linear dinucleotides such as pApA and pGpG.

**Figure 6:**
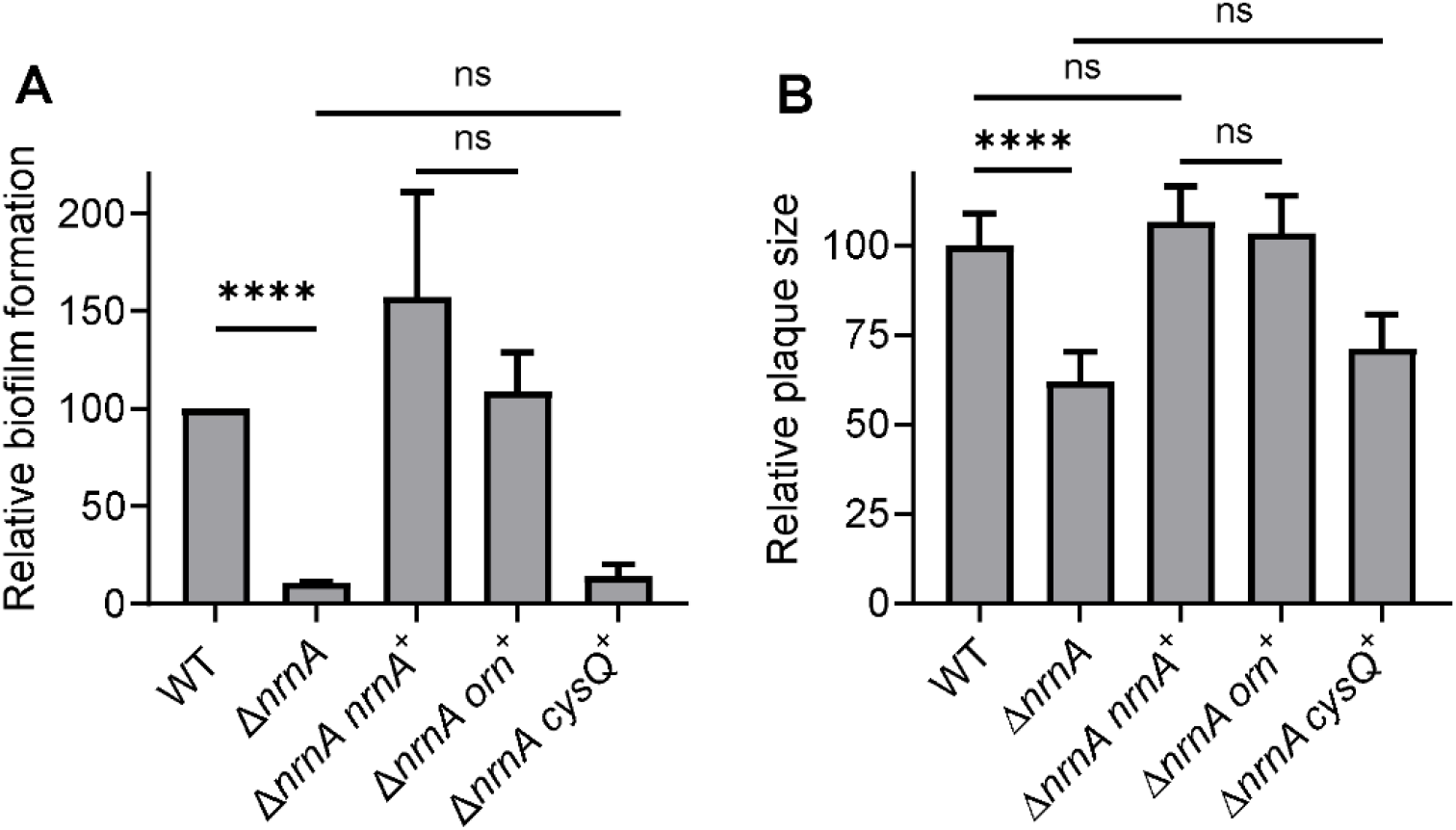
*Lm* NrnA is functionally analogous to Orn. **A** – Biofilm formation assays and **B** – Plaque formation assays, performed as for Fig. 4. The *orn* and *cysQ* genes were cloned from *E. coli* MG1655. All complementing genes were expressed under a strong constitutive promoter (P_hyper_). Data are average of at least three independent experiments, and were normalized to the WT level in each experiment. Error bars represent standard deviations. Statistical analyses were performed by paired t-tests for strains in each experiment, ns: non-significant, ****, P < 0.0001.

## DISCUSSION

By utilizing c-di-AMP and c-di-GMP signaling, *L. monocytogenes* must regulate the levels of these second messengers. The previously identified phosphodiesterases for these cyclic dinucleotides degrade them into their corresponding hydrolytic products, pApA and pGpG (21– 23). Their subsequent degradation to mononucleotides has not been examined in this bacterium. Among the reported oligoribonucleotidases, *L. monocytogenes* encodes a single homolog, NrnA. Here, we found that *Lm* NrnA robustly degrades pApA and pGpG into the mononucleotides AMP and GMP, respectively. Although the purified *Lm* NrnA enzyme can degrade c-di-AMP and c-di-GMP, these cyclic dinucleotides are unlikely physiological substrates in *L. monocytogenes*. The Δ*nrnA* mutant exhibited biofilm formation and infection defects, most likely due to the toxic accumulation of dinucleotides such as pApA and pGpG.

In the *P. aeruginosa* Δ*orn* mutant, the accumulated pGpG inhibits EAL-domain phosphodiesterases, thereby elevating c-di-GMP levels (11, 12). Similarly, the *S. aureus* Δ*pde2* mutant (analogous to *L. monocytogenes* Δ*nrnA*), accumulates both pApA and c-di-AMP (13). The accumulation of c-di-AMP might be attributed to the inhibitory effect that pApA exerts on GdpP, the major c-di-AMP phosphodiesterase in this bacterium. Additionally, this phenotype likely also reflects the hydrolytic activity of Pde2 towards c-di-AMP in *S. aureus*, since the Δ*gdpP* Δ*pde2* mutant is further elevated for c-di-AMP compared to the Δ*gdpP* strain. By contrast, *L. monocytogenes* appeared to separate cyclic dinucleotide and oligoribonucleotide metabolism. We found *Lm* NrnA to have negligible effects on c-di-AMP levels, even in the absence of both PdeA and PgpH, indicating that NrnA is specialized for oligoribonucleotide hydrolysis. Furthermore, unlike *P. aeruginosa* and *S. aureus*, the *L. monocytogenes* Δ*nrnA* mutant did not exhibit elevated c-di-AMP or c-di-GMP levels. This might reflect the absence of product inhibition on cyclic dinucleotide phosphodiesterases. Alternatively, *L. monocytogenes* might harbor additional oligoribonucleotideases, and Δ*nrnA* deletion alone might not accumulate enough pApA and pGpG for discernible inhibition of PdeA and EAL-domain phosphodiesterases in the bacterial cell, respectively. In a search for redundant phosphodiesterases, we examined *Lm* YhaM, since the *B. subtilis* YhaM homolog can modestly hydrolyze oligoribonucleotides in vitro, and weakly complement the *E. coli* and *P. aeruginosa* Δ*orn* mutants (14, 30). Although *Lm* YhaM exhibited phosphodiesterase activity, it did not degrade any tested cyclic or linear dinucleotides.

Both c-di-GMP and c-di-AMP have been shown to regulate bacterial biofilm formation. Elevated c-di-GMP levels promote the motile-sessile transition through complex regulations of the bacterial flagellar motor and extracellular polysaccharide synthesis (2). The *P. aeruginosa* Δ*orn* exhibits increased bacterial cell aggregation and exopolysaccharide production, but these phenotypes are attributed to c-di-GMP accumulation, since accumulated pGpG inhibits the EAL-domain RocR phosphodiesterase (12). The mechanisms by which c-di-AMP regulates biofilm formation are much less studied, and appear divergent among species (41–44). We found the *L. monocytogenes* Δ*nrnA* mutant to be diminished for biofilm formation. Consistent with our finding, *nrnA* gene expression is upregulated in *L. monocytogenes* biofilm (45). However, the biofilm formation defect must be independent of c-di-AMP and c-di-GMP, since the Δ*nrnA* mutant was unaltered for these cyclic dinucleotide levels compared to the WT strain. It is possible that, in the absence of NrnA, the accumulated oligoribonucleotides inhibit the expression of genes for extracellular matrix biosynthesis. Indeed, oligoribonucleotides have been shown to likely regulate the transcription of CRISPR-associated genes in *Mycobacterium* sp (10).

The *L. monocytogenes nrnA*::Himar1 mutant was previously shown to have a mouse virulence defect, and we made similar observations here using plaque formation assays in L2 cells (34). Because the Δ*nrnA* mutant was fully complemented by the *E. coli orn*^+^ allele, we hypothesize that *Lm* NrnA is functionally analogous to Orn, and therefore preferably degrades linear dinucleotides such as pApA and pGpG (15). Although short oligoribonucleotides have been shown to cause global shifts in transcriptional start sites in *P. aeruginosa* (9), we found no evidence for this to occur in *L. monocytogenes*, since the Δ*nrnA* mutant had no growth or stress response defects in broth cultures, and the expression of at least two PrfA-regulated genes, *hly* and *actA*, was intact under infection conditions. However, the Δ*nrnA* plaque defect was completely rescued upon PrfA* expression, indicating that the secretion or activity of PrfA-regulated virulence factors are impaired in this strain. Among those, the secretion of LLO (encoded by *hly*) has been shown to be unaffected in the *nrnA*::Himar1 mutant. Altogether, the regulatory functions of oligoribonucleotides in *L. monocytogenes* pathogenesis require further investigation.

## MATERIALS AND METHODS

### Bacterial strains and culture conditions

*L. monocytogenes* strains used in this study are listed in **Table S1**. *L. monocytogenes* cultures were grown in Brain Heart Infusion (BHI) broth or *Listeria* Synthetic Medium (LSM) at 37°C (19). Complementation of the Δ*nrnA* mutant was accomplished by expressing *L. monocytogenes nrnA, E. coli MG1655 orn*, and *cysQ* genes under a P_hyper_ promoter in the pPL2 plasmid, which stably integrates into the *L. monocytogenes* chromosome (46). Gene deletions were performed by allelic exchange using pKSV7 as previously described (22).

### Biofilm formation assays

Biofilm formation assays were performed for cultures grown in *Listeria* Synthetic Medium (LSM) containing 12.5 μg/mL cysteine (modified LSM) at 30°C as previously described (19, 47). Briefly, overnight cultures were inoculated into 200 μM of modified LSM in a polystyrene 96-well plate, and incubated statically for 40 hours. Total growth was assessed by measuring OD595 nm of the cultures after careful resuspension of all biomass in the wells. For biofilm quantification, culture supernatants were removed, and wells were gently washed with sterile MilliQ water. After the wells were air dried, biofilm was stained with 0.3% crystal violet (w/v), solubilized with 90% ethanol, and quantified by measuring absorbance at 595 nm.

### Protein expression and purification

The *nrnA* gene (*lmo1575*) from *L. monocytogenes* 10403S was PCR-amplified and cloned into pET20b vector with NdeI and XhoI restriction sites, generating an NrnA - 6xHis construct. The recombinant NrnA – 6xHis protein was expressed in *E. coli* Rosetta (DE3) strain and purified as previously described (48). After purification, NrnA – 6xHis was buffer exchanged using PD-10 columns (GE Healthcare) into storage buffer (40 mM Tris pH 8, 10 mM NaCl). Protein concentration was measured by Bradford assay (BioRad).

### Enzyme assays

Enzyme assays were performed in reactions containing reaction buffer (100 mM Tris pH 8, 20 mM KCl, 100 μM DTT, 2 mM MnCl_2_), NrnA, and varying substrate concentrations, incubated at 37°C. NrnA concentrations used for reactions with different substrates were: 50 nM for bis-*p*-nitrophenylphosphate (bis-*p*NPP); 1 μM for c-di-AMP and c-di-GMP; 15 nM for pAp; 0.2 nM for pApA, pGpG, and pApG. Activity towards bis-*p*NPP was monitored continuously by absorbance at 410 nm. For enzyme assays with varied divalent cations, 1 mM of each metal was used. For enzyme assays at different pH conditions, 100 mM Tris-HCl or Tris-Base was used. Reactions with nucleotide substrates were stopped at 0, 5, 10, 15, and 30 minutes by heating at 95°C for 5 minutes and analyzed on a Dionex UltiMate 3000 HPLC equipped with a DAD-3000 diode array detector. For HPLC, 20 μL of the supernatant was injected into a reverse phase column (Waters Nova-Pak C18, 3.9 x 150 mm, 4 μm) equilibrated with 100 mM potassium phosphate pH 6. Nucleotides were eluted by a mobile phase containing 100 mM potassium phosphate pH 6, over a gradient of up to 40% methanol. Concentrations of remaining substrates and formed products were calculated based on standard curves for each nucleotide.

### L2 plaque assay

Plaque assays were performed using L2 fibroblasts as previously described (49). Briefly, L2 cells were plated onto a 6-well dish at 1.2 x 10^6^ cells per well and infected with *L. monocytogenes* at an MOI of 0.5. At 1 hour post infection, 50 μg/mL gentamicin was added to cell culture media to kill extracellular bacteria. At 4 to 6 days post infection, cells were stained with 0.3% crystal violet and imaged for plaques. Areas of plaques were quantified with the ImageJ software (https://imageJ.nih.gov).

### qRT-PCR

*L. monocytogenes* cultures were grown in BHI broth at 37°C to OD600 nm of 0.7, washed, resuspended in LB broth containing 25 mM glucose-1-phosphate and 0.2% activated charcoal. Cultures were harvested immediately before and 1 hour following resuspension for RNA extraction. RNA samples were treated with Turbo DNase (Ambion, Thermo Fisher), converted to cDNA using iScript cDNA synthesis kit (BioRad), and gene expression was quantified using iTaq Universal SYBR green (BioRad) with primers specific to each target, with *rplD* as the control housekeeping gene (**Table S2**).

### Intracellular growth curves

C57BL/6 immortalized bone marrow-derived macrophages (iBMMs) were grown in DMEM with 20% heat-inactivated FBS, 2 mM sodium pyruvate, 1 mM L-glutamine, and 1% β-mercaptoethanol. Infection and intracellular growth assays were performed as previously described (50). Briefly, iBMMs were plated onto a 24-well plate at 0.4 x 10^6^ cells per well and infected with *L. monocytogenes* at an MOI of 1. At 30 minutes post infection, cells were washed with PBS and replenished with fresh medium containing 50 μg/mL gentamicin. At indicated time points, cell medium was removed, cells were lysed with MilliQ water, and serial dilutions of cell lysates were plated onto LB agar to assess bacterial burdens.

### Type I interferon assays

IFN-β production by iBMMs was detected using the type I IFN reporter cell line, ISRE-L929, as previously described (32). Briefly, ISRE-L929 cells were grown in DMEM with 5% heat-inactivated FBS and plated onto a 96-well plate at 5 x 10^5^ cells per well. iBMMs were infected with *L. monocytogenes* as described for intracellular growth curves. At 5 hours post infection, iBMM cell culture supernatants were removed and incubated with ISRE-L292 cells for 4 hours. Uninfected iBMM supernatant was used as background control. Following incubation, L929 cells were lysed with 40 μL TNT buffer (20 mM Tris-Base, pH 8, 100 mM NaCl, 1% Triton X-100). Finally, L292 cell lysates were mixed with 40 μL of luciferase substrate solution (20 mM Tricine, 2.67 mM MgSO4.7H2O, 0.1 mM EDTA, 33.3 mM DTT, 530 μM ATP, 270 µM acetyl CoA lithium salt, 470 μM luciferin, 5 mM NaOH, 265 µM magnesium carbonate hydroxide). Bioluminescence was measured using a Synergy H1 plate reader (BioTek).

### Quantification of nucleotides by LC-MS/MS

Quantification of c-di-AMP and c-di-GMP from *L. monocytogenes* cultures were performed as previously described (22). Briefly, *L. monocytogenes* strains were grown in BHI to mid-log phase (OD600 nm ∼0.5). Bacterial cell pellets from 0.5mL cultures were resuspended in 50µL of 0.25µM C^13^N^15^-c-di-AMP, used as an internal standard, and lysed by sonication. Fractions from methanol extraction were pooled, dried by evaporation, and resuspended in 50 μL of MilliQ H_2_O. LC-MS/MS was performed on an Agilent 6460 Triple Quad LC-MS/MS with a 1260 HPLC. Chromatographic separation was performed with an analytical Synergi 4u Hydro-RP 80A column (50 × 2 mm, 4 μm, Phenomenex). Mass transitions for c-di-AMP quantification were previously published (22). C-di-GMP was detected based on three transitions: +691/152, +691/540, and +691/248, and quantified using the +691/152 transition. For each nucleotide, the peak areas of the quantifier transition were divided by that of C^13^N^15^-c-di-AMP (+689/146) in the same sample.

## ACKNOWLEDGEMENTS

*L. monocytogenes* strains Δ*hly* and Δ*nrnA* are generous gifts from Dr. John-Demian Sauer. We thank Dr. Brad Bolling for access to the HPLC and technical assistance with the instrument. A.R.G. was funded by a postdoctoral fellowship from the Wisconsin Dairy Innovation Hub. T.N.H was funded by the Foundation for Food and Agricultural Research and a USDA Hatch Act Formula Fund.

